# Inference of significant microbial interactions from longitudinal metagenomics sequencing data

**DOI:** 10.1101/305326

**Authors:** Xuefeng Gao, Bich-Tram Huynh, Didier Guillemot, Philippe Glaser, Lulla Opatowski

## Abstract

Data of next-generation sequencing (NGS) and their analysis have been facilitating advances in our understanding of microbial ecosystems such as human gut microbiota. However, inference of microbial interactions occurring within an ecosystem is still a challenge mainly due to metagenomics sequencing (e.g., 16S rDNA sequences) providing relative abundance of microbes instead of absolute cell count. In order to describe the population dynamics in microbial communities and estimate the involved microbial interactions, we introduce a procedure by integrating generalized Lotka-Volterra equations, forward stepwise regression and bootstrap aggregation. First, we successfully identify experimentally confirmed microbial interactions with relative abundance data of a cheese microbial community. Then, we apply the procedure to time-series of 16S rDNA sequences of gut microbiomes of children who were progressing to Type 1 diabetes (T1D progressors), and compare their gut microbial interactions to a healthy control group. Our results suggest that the number of inferred microbial interactions increased over time during the first three years of life. More microbial interactions are found in the gut flora of healthy children than the T1D progressors. The inhibitory effects from *Actinobacteria* and *Bacilli* to *Bacteroidia*, from *Bacteroidia* to *Clostridia*, and the benifit effect from *Clostridia* to *Bacteroidia* are shared between healthy children and T1D progressors. An inhibition of *Clostridia* by *Gammaproteobacteria* is found in healthy children that maintains through their first three years of life. This suppression appears in T1D progressors during the first year of life, which transforms to a commensalism relationship at the age of three years old. *Gammaproteobacteria* is found exerting an inhibition on *Bacteroidia* in the T1D progressors, which is not identified in the healthy controls.

## Background

The human gut microbiota, an ecosystem comprising a diverse collection of interactive microbial species, plays a key role in nutrition, metabolism, physiology and immune function in humans [1]. An interaction between two microbes can either be neutral (0), positive (+) or negative (−) to the fitness of the interacting microbes. Based on the overall effects, an interaction can be classified into commensalism (0/+), amensalism (0/−), mutualism (+/+), parasitism (+/−), competition (−/−) or no interactions (0/0). Interactions among the microbial species characterize the composition and function of gut microbiota, thereby influencing the host’s health [2]. The microbial interactions in the gastrointestinal tract are complex and flexible, capable of adapting to physiological perturbations [3]. The change of one species may shift the relative abundances of other members in the community and affect the community’s functional capacity. A better understanding of ecological dynamics and microbial interactions is essential to investigate the consequences of taxonomic perturbations.

By using metagenomics sequencing technologies, such as 16S rRNA gene profiling, it is now possible to follow the time-evolution of a microbial population by measuring abundance of bacterial species in a community. However, analyzing the dynamics of bacterial community from genomic survey data is not straightforward, because metagenomics sequencing provides relative abundances of microbial species based on a fixed total number of sequences rather than absolute cell count. The high density of genomic survey data and its compositional nature are the most common challenges involved in inferring microbial interaction networks. Hence, the power of computational approaches allows to address this challenge. Mathematical and computational approaches make it possible to analyze highly complex microbial communities.

Correlation networks have proved useful for detecting biological interactions. There are many different techniques for computing correlation networks, for example Pearson correlation coefficient [4], Spearman correlation coefficient [5], Bray-Curtis distance [6], Local Similarity Analysis [7–10], Maximal Information Coefficient [11], SparCC [12], based on Aitchison’s log-ratio analysis [13] and CoNet which combine information from several different standard comparison metrics [14]. Weiss *et al*. [15] benchmarked the performance of these techniques in dealing with the relative abundance in simulated data specific to microbiome studies. Despite the correlation networks are useful in studying some overall biological relationships, they have significant limits in deciphering non-linear interactions such as between three or more species, and are unable to detect relationships like amensalism and partial-obligate-syntrophy [15,16]. Indeed, two species may be correlated even if they do not directly interact with each other.

The use of non-linear differential equations like Lotka-Volterra (LV) model is an alternative approach to study microbial interactions. The generalized Lotka-Volterra (gLV) equations are able to describe the time-dependent population dynamics and predict ecological relationships (i.e. mutualism, commensalism, parasitism and competition) between members of different biological species. Several studies have attempted to use gLV equations studying microbial communities that consist of multiple bacterial species [17–20]. In particular, Mounier et al. used gLV equations describing the population dynamics of a cheese community consisting of five microbial groups with experimental data of the absolute cell numbers [21]. This model predicted some microbial interactions that were confirmed by co-culture experiments afterwards.

Fisher and Mehta [18] have proposed an approach named Learning Interactions from Microbial Time Series (LIMITS), which implements forward stepwise regression with median bootstrap aggregation for gLV equations inferring the species interactions in the human gut microbiomes. By combining forward stepwise regression with bootstrap aggregation, LIMITS was able to overcome a statistical issue termed “errors-in-variables” (accounting for measurement errors in the independent variables) and infer species interactions from time series relative abundances of species [22]. However, these inferred interactions were not experimentally confirmed.

Although these tools brought some insights to infer microbial interactions, they have no proven ability to infer biologically confirmed microbial interactions from relative abundance data. Here, we propose a procedure by integrating gLV equations, forward stepwise regression, bootstrap aggregation and one-sample t-test. First, we test and validate this procedure by correctly identifying microbial interactions from the relative abundances of a cheese microbial community [21]. For application, we then apply our procedure to infer microbial interactions within gut microbiota of healthy and type 1 diabetes (T1D) progressing infants (seroconverters for diabetes autoimmune) from a set of reported data [23].

## Results

### Procedure validation on experimentally confirmed data: identification of significant microbial interactions within a cheese microbial community

The procedure was tested through describing the dynamics of relative abundances of five microbial groups and estimating their interactions within a cheese microbial community. For initial conditions, the intrinsic growth rates and inter-species interaction coefficients were set at 1 generation/day and 0, respectively. The carrying capacity of the cheese microbial community is 10^10^ CFU/g [21].

The gLV modeling succeeded in describing the changes of relative abundance of the microbial populations, and roughly predicted the growth trends of each member of this community (Figure 2). The inferred intrinsic growth rates of the microbes and their significant interactions were shown in Figure 3. In particular, some of the inferred interactions (Figure 3B) were previously confirmed in the co-culture experiments, including the promotion effects from *G. candidum* to the bacterial group and *Leucobacter sp*., and the inhibitory effects of *G. candidum* on *D. hansenii* [21]. The estimated intrinsic growth rate of the bacterial group was low (Figure 3A), indicating that their growth was highly influenced by other members in the community. A benefit effect of *G. candidum* on the bacterial group was inferred, suggesting that the growth of the bacterial group partially relies on *G. candidum*. According to a co-culture study [21], the growth of some bacteria, such as *Brevibacterium aurantiacum*, was actually relying on *G. candidum* was identified. An inhibitory effect of *D. hansenii* on the bacterial group was predicted by our procedure (Figure 3B). This interaction may be explained by the antagonistic activity of some *D. hansenii* strains against bacteria (via production of antagonistic metabolites) [24].

**Figure 1.**
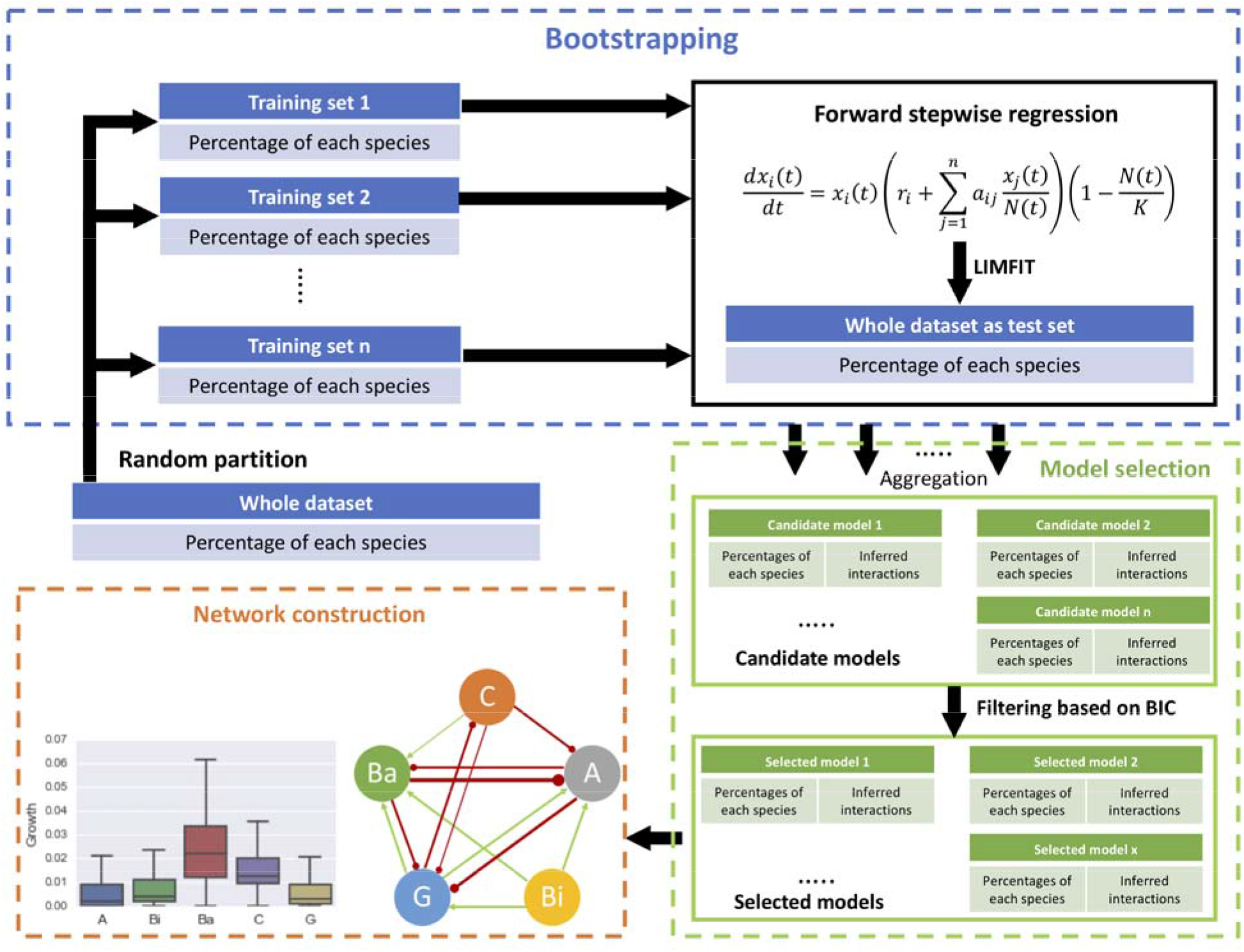
Schematic diagram of the proposed procedure, consisting of forward stepwise regression, bootstrap aggregating, model selection, and network construction.

**Figure 2.**
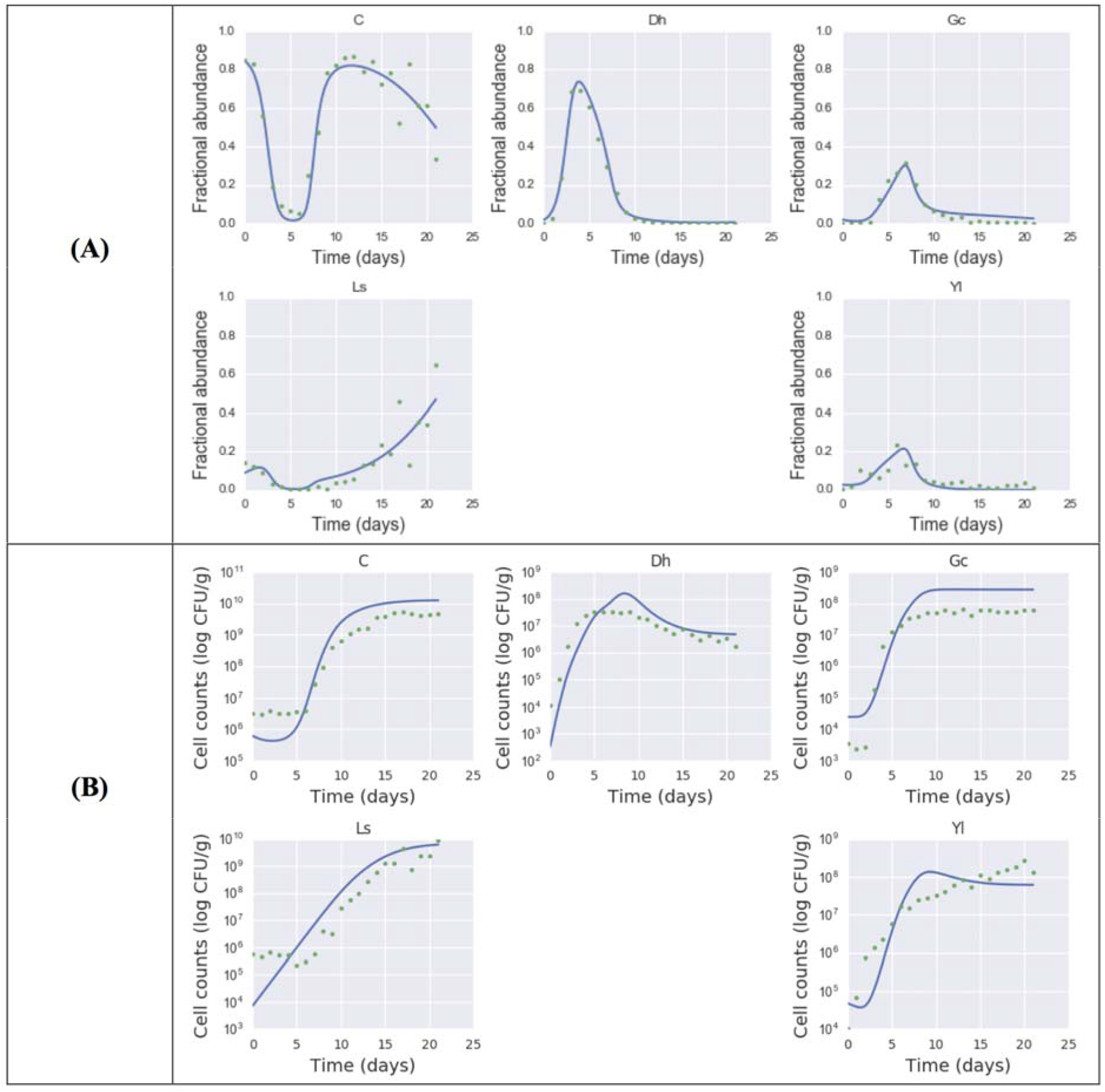
An example of using gLV equations and forward stepwise regression describing the population dynamics of a multispecies ecosystem in **(A)** proportions and **(B)** cell counts. Experimental data [21] is indicated by dots and the predicted population dynamics are indicated by curves. Abbreviations: A: *Actinobacteria*; Bi: *Bacilli*; Ba: *Bacteroidia*; C: *Clostridia*; G: *Gammaproteobacteria*.

**Figure 3.**
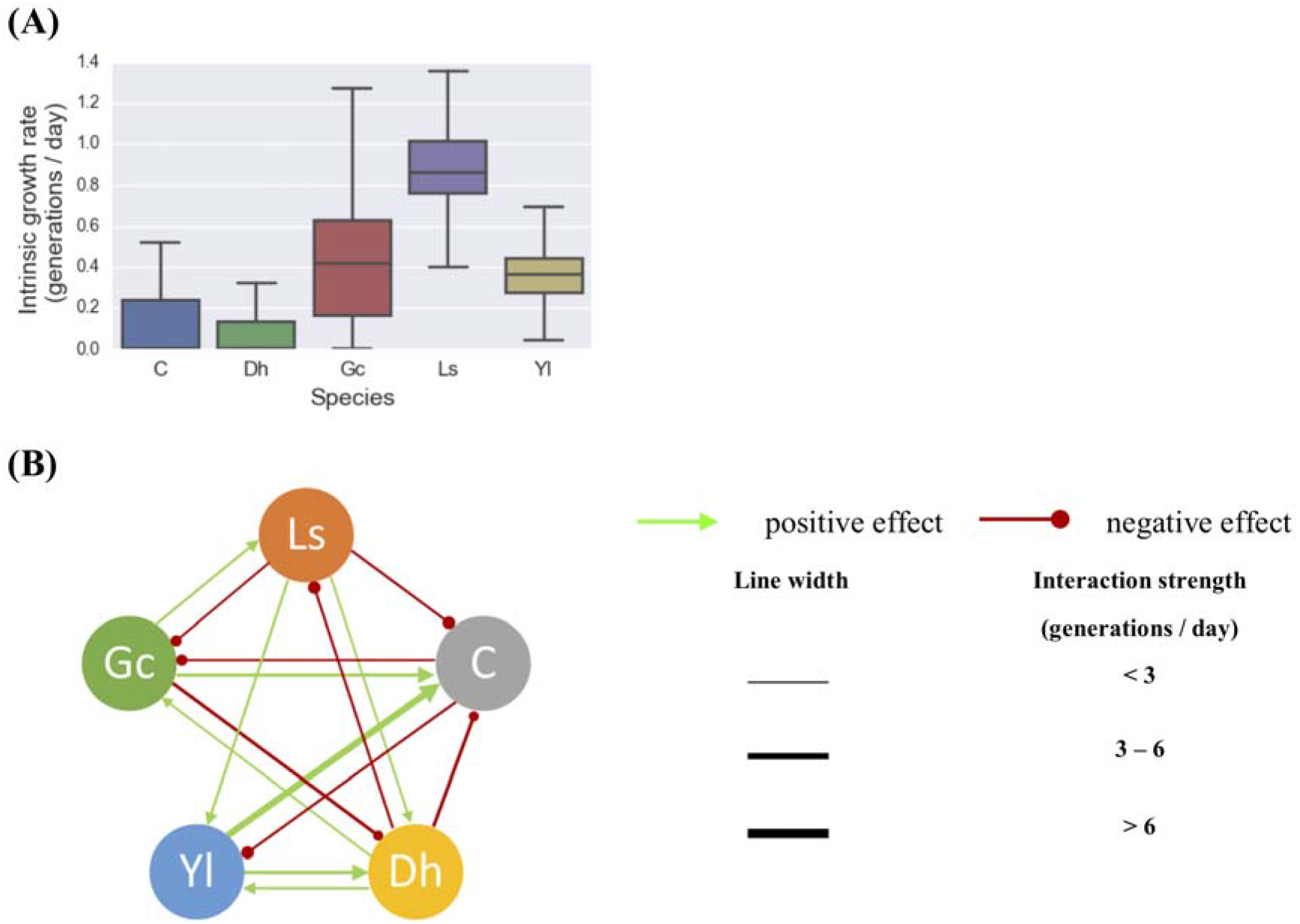
Inferring significant microbial interactions within a cheese microbial community. **(A)** The intrinsic growth rates of the microbes and **(B)** their significant interactions were inferred from the longitudinal data of five microbes. 80% of the whole data set was randomly selected for training the gLV model, with 1000 bootstrap samples. Resulted interactions were selected with one-sample t-test *P*(*a*_ij_ ≠ 0) > 95%. The line thickness is proportional to the strength of the interaction. Abbreviations: Dh: *D. Hansenii*; Yl: *Y Lipolytica*; Gc: *G. Candidum*; Ls: *Leucobacter sp*.; C: *Arthrobacter arilaitensis, Hafnia alvei, Corynebacterium casei, Brevibacterium aurantiacum, and Staphylococcus xylosus*.

The sensitivity of our procedure was assessed by varying the number of bootstrapping samples and the size of training data. The performance of this procedure was influenced by the number of bootstrap samples. We observed that some significant interactions were detected using a relatively low number of bootstrap samples (e.g., *D. hansenii* inhibiting the bacterial group and *G. candidum* promoting the bacterial group) while some others appeared along with increased number of bootstraps (Table S1). The frequency of these interactions may be an indicator of their weight in shaping the community dynamics.

In summary, our procedure was able to identify ground-truth microbe-microbe interactions within a microbial community through relative abundance data.

### Application on time-series 16S rRNA gene sequence data: gut microbial interaction networks of healthy children versus T1D progressors

Next, we used this approach to study the gut microbial dynamics in healthy children and those who were progressing to T1D during their first three years of life. Kostic et al. [23] examined the composition dynamics of gut microbiomes in 33 children genetically predisposed to T1D from birth until three years of age with monthly sampling. The authors observed a relative reduction in alpha-diversity in the gut of children who progress to T1D compared to the seroconverters defined as positive for at least two autoantibodies (no T1D developers occurred during the follow-up) and in non-converters’ gut. We applied the proposed procedure on the 16S rRNA sequencing data reported in [23]. We investigated the class level gut bacterial interactions in T1D-associated and healthy infants at different age stages. The carrying capacity of intestinal microbial community was assumed to be 10^10^ cells/g of feces [25]. The resulting intrinsic growth rates and interaction coefficients of the five bacterial classes were given in Table S2 for the two groups at different age stages.

The gut microbial interaction network consisted of just a few amensalism interactions for the first year of life, but became more complex over time (Figure 4). There were more bacterial interactions identified in healthy children than T1D progressors after the age of one year old. Some interactions were shared between healthy controls and T1D progressors, including the inhibitory effects *Gammaproteobacteria* to *Clostridia* for one year of age, the inhibitory effect from *Actinobacteria* to *Bacteroidia* and the promotion effect from *Clostridia* to *Bacteroidia*.

**Figure 4.**
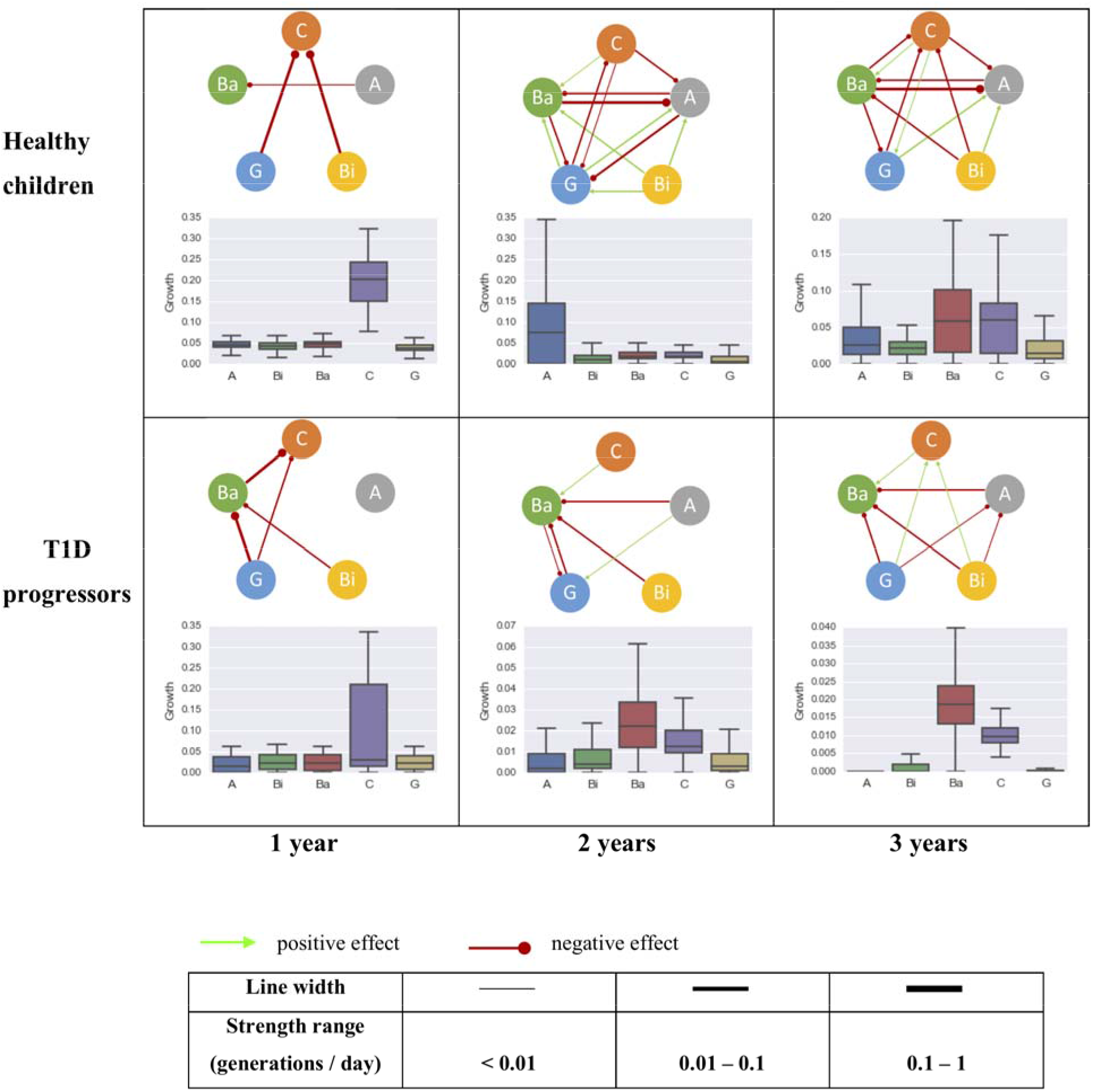
The intrinsic growth rates of five most abundant bacterial classes and their interactions within the gut microbiome community of healthy (top row, control) and T1D progressors (bottom row, case) at 0-1 year (left column), 0-2 years (mid column) and 0-3 years (right column) of age. The line thickness is proportional to the strength of the interaction. Abbreviations: A: *Actinobacteria*; Bi: *Bacilli*; Ba: *Bacteroidia*; C: *Clostridia*; G: *Gammaproteobacteria*.

Three interactions were identified uniquely in the healthy controls: inhibitions of *Actinobacteria* by *Bacteroidia* and *Clostridia*, and a promotion from *Gammaproteobacteria* to *Actinobacteria*. An inhibitory effect from *Gammaproteobacteria* to *Clostridia* was established within the first year of life which maintained through the following three years (top row of Figure 4). Intriguingly, the effect from *Gammaproteobacteria* to *Clostridia* was identified as amensalism during the first year of life for T1D progressors, which transformed to a benefit effect at their age of three. An inhibitory effect from *Gammaproteobacteria* to *Bacteroidia* was only observed in the T1D progression group (bottom row of Figure 4). The relationship between *Bacilli* and *Clostridia* was amensalism for healthy children, but commensalism for T1D progression children at age of three.

## Discussions

Investigating the gut microbial community composition and their interaction patterns are crucial to understand the role of gut microbiota in maintaining human health and causing diseases. However, a systematic approach for accessing microbial interactions has been lacking. Some methods have been attempted to infer microbial interaction networks, such as correlation networks and mathematical modeling. Although these attempts have contributed in creating interaction networks, none of their resulting microbial interactions have been experimentally confirmed. In the present study, we showed that a combination of gLV equations, forward stepwise regression, and bootstrap aggregation was able to infer microbial interactions from longitudinal relative abundance data. This proposed procedure was able to infer experimentally confirmed microbial interactions within a cheese microbial community[21], such as the promotions of *G. candidum* on *Leucobacter sp*. and on a group of bacteria, and a suppression of *D. hansenii* by *G. candidum*. In addition, the estimated changes in cell counts by our procedure roughly recapitulated the growth trends of each microbial group in the community. To our knowledge, this is the first study showing experimentally confirmed interactions inferred from relative abundance of taxa within a microbial community. Despite the potential of our proposed method in dealing with compositional data, it could not explain all the phenomena in the coculture experiments. For example, our procedure predicted a relative higher intrinsic growth rate for *Leucobacter sp*‥ Indeed, the growth of *Leucobacter sp*. was found to depend on *G. candidum* [21] and had a long lag phase at the start of ripening, which were not observed from the results of our procedure.

One goal of comparative metagenomics is to identify meaningful changes in the microbiome’s taxonomic and functional composition that are associated with health and disease. Disruptions of the process of gut microbial colonization during childhood have been shown to be associated with an increased risk of adiposity and pathogenesis of autoimmune disorders such as T1D [23,26–28]. To address how alterations of the gut microbial community composition may contribute to childhood disease, we must first investigate the normal dynamics of the community in the growing infant. By comparing the gut taxonomic trajectories of children who progress to T1D compared to healthy controls, Kostic et al. [23] have found general changes in abundance of the gut microbial community and the timing of these changes. They identified some microbes strongly correlated which was consistent across most healthy subjects ([23], such as a positive correlation between *Gammaproteobacteria* and *Actinobacteria* or a negative correlation between *Gammaproteobacteria* and *Clostridia* [23]. These correlations can be explained by the inferred interactions from our procedure, e.g., promotion effect from *Gammaproteobacteria* to *Actinobacteria* and inhibition effect from *Gammaproteobacteria* to *Clostridia*.

By applying our procedure to a longitudinal time-series study of gut microbiota in children [23], we demonstrated that the gut bacterial interaction network (at class level) was getting more complex along with age, with more interactions inferred for the healthy children than the T1D progressors. Some microbial interactions were shared between healthy children and the T1D progressors, such as inhibitory effects from *Actinobacteria* and *Bacilli* to *Bacteroidia*. Although the composition of the microbiome community was changing along with child development, some of the bacterial interactions, in particular at higher phylogenetic levels such as class, remained stable after establishment. For example, the inhibitory effect from *Gammaproteobacteria* to *Clostridia* was established for healthy children cohort in the first year of life and maintained through the age of three. In T1D progressors, whereas the amensalism relationship between *Gammaproteobacteria* and *Clostridia* appeared during their first year of life, which became commensalism (i.e., *Gammaproteobacteria* enhancing the growth of *Clostridia*) at the age of three. The inhibition from *Gammaproteobacteria* to *Bacteroidia* maintained during first three years of life for T1D progressors, which was not detected in the healthy controls.

It has been observed that the quantity of *Clostridium* (genus level, belongs to the class *Clostridia*) correlated positively and significantly with the plasma glucose level in diabetic children [26]. In newborn infants, *Gammaproteobacteria* have been shown to cause a healthy level of inflammation in their intestines, protecting them from excessive inflammatory and autoimmune disorders later in life [29]. From the data of [23], the proportion of *Gammaproteobacteria* was indeed lower in the T1D progression children group than healthy controls during the first months of life (e.g., about 11% vs. 16% for the two groups within 180 days after birth). In addition, *Bacilli* was also found to promote the growth of *Clostridia* in the gut community of T1D progression children, whereas it contributed to inhibition of *Clostridia* for the healthy controls. Thus, our results suggest that the inhibitory effects of *Gammaproteobacteria* and *Bacilli* on *Clostridia* might play a role in regulating plasma glucose in early lives and protect children from developing to T1D. Further study is necessary to test this hypothesis.

## Conclusions

Here we propose a procedure which was capable of inferring experimentally confirmed microbial interactions out of compositional data. By using our procedure, we identified some similarities as well as differences in the gut bacterial interaction networks between children towards T1D progression and healthy controls in their first three years of life. The interaction networks were getting more complex in the gut microbiome of children after their first year of life. The number of interactions was higher in healthy children than in T1D progressors. Some bacterial interactions were exclusively found in each group, which may be able to predict the T1D progression state. Our results provide potential new insights into the relationship between gut microbial interactions and infants’ T1D progression. In the future, by incorporating microbiome’s functional shift [30], the procedure presented above might help to interpret disease-specific changes of microbial interactions and accordingly help to predict disease progression.

## Methods

A diagram of our procedure was given in Figure 1.

### Generalized Lotka-Volterra equations

We used gLV equations to describe an ecological community consisting of *n* microbiome taxa. For taxon *i*, the population dynamics is described as

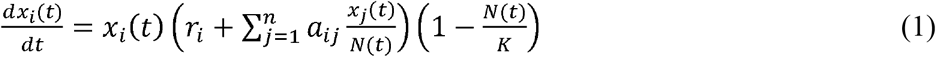

where *x*_*i*_(*t*) is the abundance of taxon *i* at time *t, r*_i_ the intrinsic growth rate of taxon *i*, and *a*_ij_ the effect that taxon *j* has upon taxon *i*, which is proportional to the proportion of species *j*. We assumed the interactions between species *i* and *j* is not necessarily bilateral, which can take one of six possible forms based on the signs of *a*_ij_ / *a*_ji_ (*i* ≠ *j*); +/- (parasitism), -/- (competition), +/+ (mutualism), +/0 (commensalism), -/0 (amensalism) and 0/0 (neutral). *N*(*t*) represents the community size at time *t* (i.e., 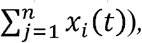, and *K* the carrying capacity of the community (a logistic growth component). For simplicity, we did not consider intra-species interactions, such as the competition for resources between the members of a same species. However, a carrying capacity of the environment was introduced. We fitted relative abundance data to the gLV equations using the *Levenberg-Marquardt* algorithm for non-linear least squares minimization, implemented by lmfit [31] which is a high-level interface to non-linear optimization and curve fitting tool for Python.

### Forward stepwise regression

We used a forward stepwise regression method for the gLV model selecting interactions that explain most of the population dynamics of microbial species. This method started with no interaction coefficients in the model (i.e., only with growth terms), testing the addition of each interaction coefficient using a comparison criterion, adding the interaction if it improves the model the most, and repeating this process until no improvement of the model fitting. Stepwise regression is prone to overfitting the data, which arises from searching a large space of possible models. We use Bayesian information criterion (BIC) [32] as a selection criterion in order to solve the overfitting problem by introducing a penalty term for the number of parameters in the model. BIC is calculated by

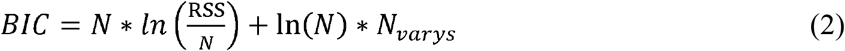

where, *RSS* is the residual sum of squares over the whole data set, *N* is the number of data points and *N*_varys_ the number of variable parameters in the model. Therefore, a lower BIC implies either a better fit, fewer explanatory variables, or both.

### Bootstrap aggregating (Bagging)

In order to attenuate the instability problem of forward stepwise regression, we introduced bootstrap aggregating strategy for noise-reduction [33] by randomly partitioning the data into two sets: training set and test set. A typical way to test for accuracy in models created by stepwise regression is to evaluate the models against a test set that was not used for model creation. Accuracy is then calculated as errors between the values predicted by the model and data in the test set. In order to assess both the descriptive and predictive power, we fitted the models to the training set but evaluate the models to the whole data set. The procedure is performed in the following steps:

1. At each iteration, partitioning the data into a training set and a test set.
2. The gLV model fitting on the training set and BIC calculated for the best parameters starting with only growth terms, and all interaction coefficients are assigned to a parameter set TEST{*a*_ji_}.
3. Each single and pairwise interaction coefficient in the TEST{*a*_ji_} is separately added to current model, then a BIC is calculated over the whole data set. After scanning over all coefficients in the TEST{*a*_ji_}, the one (or pairwise) that generated the lowest BIC is transferred to a parameter set SELECT{*a*_ji_}. If multiple models have similar BIC values that are close to the lowest BIC (ΔBIC < 2), one of them is randomly selected. The new model then consists of all coefficients in SELECT{*a*_ji_}.
4. Repeating step 3 for the new model to evaluate the model fitting after addition of (remaining) coefficients in the TEST{*a*_ji_} and update the lowest BIC accordingly.
5. Iteration stops when additional coefficients no longer reduce the lowest BIC. The resulted coefficients in the SELECT{*a*_ji_} comprise an estimate of the interaction network.

This forward stepwise regression process was repeated many times (up to 1000 bootstraps in this study), each resulting in a set of interaction coefficients that compose a specific gLV model. These models were then aggregated. Next, we removed models that violated biological constrains (e.g., based on a minimum population size and/or a minimum number for each species in the community). After filtering, we selected models with minimum BIC values (the minimum BIC + 10% allowance), and their parameters (growth rates and interaction coefficients) were compiled. Finally, the significant interactions were determined through one-sample t-test with the value of *P*(*a*_ij_ ≠ 0) > 95%.

### Data

Mounier et al. [21] investigated interactions between yeast and bacteria within a model choose composed of three yeasts (Debaryomyces hansenii 1L25, Geotrichum candidum 3E17, and Yarrowia lipolytica 1E07) and six bacteria (Arthrobacter arilaitensis 3M03, Brevibacterium aurantiacum 2M23, Corynebacterium casei2M01, Hafnia alvei 2E12, Leucobacter sp. strain 1L36, and Staphylococcus xylosus 1L18). Two experiments were conducted to measure the microbial dynamics during the development of the ecosystem. In the first one, the dynamic of the full ecosystem was studied. Cheeses were sampled in duplicate every day for 21 days for microbial enumeration in together with measurements of lactose, lactate content, and pH. The second experiment aimed to investigate the effect of the absence of one, two or three yeasts (all combination were tested) in the ecosystem development. To test our procedure, we studied the development dynamics of this cheese microbiome community (data were obtained from [21]). Kostic et al. studied the impact of the gut microbiome dynamic on type 1 diabetes (T1D). They recruited between September 2008 and August 2010 in Finland and Estonia 33 newborns with positive cord blood testing for HLA DR-DQ, which are alleles conferring risk of T1D. The infants were followed-up until the age of 3 years old with monthly stool samples. Data regarding infections, drugs consumption (in particular antibiotics), diet were collected. Serum samples were collected at 0 (cord blood), 3, 6, 12, 18, 24, and 36 months to test for 4 diabetes-associated autoantibodies. 16S rRNA sequencing were performed for accessing the fecal microbiota composition. Eleven children had at least two positive autoantibodies seroconversion during their follow-up but did not develop T1D (i.e., seroconverters). Four children progressed to T1D by end of the study. The 11 seroconverters were matched with the 22 healthy controls for gender, HLA genotype and country. We applied our procedure on a set of 16S rDNA sequences (in OTUs) of the gut microbiota of 11 T1D progression children (seven seroconverters and four who progressed to T1D) and 22 healthy controls between the age of 0-3 years old (data were obtained from Table S2 published in [23]). In specific, we used the relative abundance of five bacterial classes: *Actinobacteria; Bacteroidia, Bacilli, Clostridia and Gammaproteobacteria*.

## Competing interests

The authors declare no competing financial interests.

## Funding

This work was supported by the program “Projets Transversaux de Recherche” PTR 2015 (grant n° PTR558). This work was also supported directly by internal resources of the French National Institute for Health and Medical Research (Inserm), the Institut Pasteur, the University of Versailles-Saint-Quentin-en-Yvelines (UVSQ) and the French Government’s “Investissement d’Avenir” program, Laboratoire d’Excellence “Integrative Biology of Emerging Infectious Diseases” (grant n°ANR-10-LABX-62-IBEID).

## Author contributions

X.G. and L.O. conceived the method and analyzed the data. X.G, L.O. wrote the manuscript with help from B.H, D.G, and P.G.

---

**Table S1**. Sensitivity analysis of the number of bootstrap samples and the size of training dataset. The strengths of significant interactions within a cheese microbial community[21], resulted from 100 - 1000 bootstraps with one-sample t-test *P*(*a*_ij_ ≠ 0) > 95%. 50%-90% of the whole data set was randomly selected for training the gLV model.

**Table S2**. The strengths of significant gut microbial interactions (class level) for the health children and the T1D progressors [23], resulted from 1000 bootstraps with one-sample t-test *P*(*a*_ij_ ≠ 0) > 95%. 80% of the whole data set was randomly selected for training the gLV model.

## References

1. Sekirov I, Russell SL, Antunes LCM, Finlay BB. Gut Microbiota in Health and Disease. Physiol. Rev. 2010;90.

2. Ríos-Covián D, Ruas-Madiedo P, Margolles A, Gueimonde M, de Los Reyes-Gavilán CG, Salazar N. Intestinal Short Chain Fatty Acids and their Link with Diet and Human Health. Front. Microbiol. 2016;7:185.

3. Sun J, Chang EB. Exploring gut microbes in human health and disease: Pushing the envelope. Genes Dis. 2014;1:132-9.

4. Pearson K. Determination of the coefficient of correlation. Science. 1909;30:p.23-25.

5. Spearman C. The Proof and Measurement of Association between Two Things. Am. J. Psychol. 1904;15:72.

6. Bray JR, Curtis JT. An Ordination of the Upland Forest Communities of Southern Wisconsin. Ecol. Monogr. 1957;27:325-49.

7. Ruan Q, Dutta D, Schwalbach MS, Steele JA, Fuhrman JA, Sun F. Local similarity analysis reveals unique associations among marine bacterioplankton species and environmental factors. Bioinformatics. 2006;22:2532-8.

8. Steele JA, Countway PD, Xia L, Vigil PD, Beman JM, Kim DY, et al. Marine bacterial, archaeal and protistan association networks reveal ecological linkages. ISME J. 2011;5:1414-25.

9. Beman JM, Steele JA, Fuhrman JA. Co-occurrence patterns for abundant marine archaeal and bacterial lineages in the deep chlorophyll maximum of coastal California. ISME J. 2011;5:1077-85.

10. Xia LC, Ai D, Cram J, Fuhrman JA, Sun F. Efficient statistical significance approximation for local similarity analysis of high-throughput time series data. Bioinformatics. 2013; 29:230-7.

11. Reshef DN, Reshef YA, Finucane HK, Grossman SR, McVean G, Turnbaugh PJ, et al. Detecting novel associations in large data sets. Science. 2011;334:1518-24.

12. Friedman J, Alm EJ. Inferring correlation networks from genomic survey data. PLoS Comput. Biol. 2012;8:e1002687.

13. Anders S, Huber W. Differential expression analysis for sequence count data. Genome Biol. 2010;11:R106.

14. Faust K, Sathirapongsasuti JF, Izard J, Segata N, Gevers D, Raes J, et al. Microbial cooccurrence relationships in the Human Microbiome. Ouzounis CA, editor. PLoS Comput. Biol. 2012;8:e1002606.

15. Weiss S, Van Treuren W, Lozupone C, Faust K, Friedman J, Deng Y, et al. Correlation detection strategies in microbial data sets vary widely in sensitivity and precision. Isme J. 2016;10:1-13.

16. Morris BEL, Henneberger R, Huber H, Moissl-Eichinger C. Microbial syntrophy: interaction for the common good. FEMS Microbiol. Rev. 2013;37:384-406.

17. Stein RR, Bucci V, Toussaint NC, Buffie CG, Rätsch G, Pamer EG, et al. Ecological Modeling from Time-Series Inference: Insight into Dynamics and Stability of Intestinal Microbiota. PLoS Comput. Biol. 2013;9:31-6.

18. Fisher CK, Mehta P. Identifying keystone species in the human gut microbiome from metagenomic timeseries using sparse linear regression. PLoS One. 2014;9:e102451.

19. Marino S, Baxter NT, Huffnagle GB, Petrosino JF, Schloss PD. Mathematical modeling of primary succession of murine intestinal microbiota. Proc. Natl. Acad. Sci. U. S. A. 2014;111:439-44.

20. Bucci V, Tzen B, Li N, Simmons M, Tanoue T, Bogart E, et al. MDSINE: Microbial Dynamical Systems INference Engine for microbiome time-series analyses. Genome Biol. 2016;17:121.

21. Mounier J, Monnet C, Vallaeys T, Arditi R, Sarthou AS, Hélias A, et al. Microbial interactions within a cheese microbial community. Appl. Environ. Microbiol. 2008;74:172-81.

22. Fuller WA. Properties of Some Estimators for the Errors-in-Variables Model. Ann. Stat‥ Institute of Mathematical Statistics; 1980;8:407-22.

23. Kostic ADD, Gevers D, Siljander H, Vatanen T, Hyötyläinen T, Hämäläinen A-M, et al. The Dynamics of the Human Infant Gut Microbiome in Development and in Progression toward Type 1 Diabetes. Cell Host Microbe. 2015;17:260-73.

24. Liu S-Q, Tsao M. Biocontrol of dairy moulds by antagonistic dairy yeast Debaryomyces hansenii in yoghurt and cheese at elevated temperatures. Food Control. 2009; 20 (9): 852-855.

25. Savino F, Quartieri A, De Marco A, Garro M, Amaretti A, Raimondi S, et al. Comparison of formula-fed infants with and without colic revealed significant differences in total bacteria, Enterobacteriaceae and faecal ammonia. Acta Paediatr. 2016; 106: 573-578.

26. Murri M, Leiva I, Gomez-Zumaquero JM, Tinahones FJ, Cardona F, Soriguer F, et al. Gut microbiota in children with type 1 diabetes differs from that in healthy children: a case-control study. BMC Med. 2013;11:46.

27. Blustein J, Attina T, Liu M, Ryan AM, Cox LM, Blaser MJ, et al. Association of caesarean delivery with child adiposity from age 6 weeks to 15 years. Int. J. Obes. (Lond). 2013;37:900-6.

28. Huh SY, Rifas-Shiman SL, Zera CA, Edwards JWR, Oken E, Weiss ST, et al. Delivery by caesarean section and risk of obesity in preschool age children: a prospective cohort study. Arch. Dis. Child. 2012;97:610-6.

29. Mirpuri J, Raetz M, Sturge CR, Wilhelm CL, Benson A, Savani RC, et al. Proteobacteria-specific IgA regulates maturation of the intestinal microbiota. Gut Microbes. Taylor & Francis; 2014;5:28-39.

30. Manor O, Borenstein E, Abubucker S, Segata N, Goll J, Schubert AM, et al. Systematic Characterization and Analysis of the Taxonomic Drivers of Functional Shifts in the Human Microbiome. Cell Host Microbe. 2017;21:254-67.

31. Newville, Stensitzki, Allen, Ingargiola. LMFIT: Non-Linear Least-Square Minimization and Curve-Fitting for Python¶. http://cars9.uchicago.edu/software/python/lmfit/confidence.html.

32. Kass RE, Wasserman L. A Reference Bayesian Test for Nested Hypotheses and its Relationship to the Schwarz Criterion. J. Am. Stat. Assoc. 1995; 90: 928-34.

33. Breiman L. Bagging Predictors. Breiman, L. Mach Learn. 1996;24:p.123-40.

